## Supplementary Material for "SERBP1 interacts with PARP1 and is present in PARylation-dependent protein complexes regulating splicing, cell division, and ribosome biogenesis"

^†^ Joint Authors

**SUPPLEMENTARY MATERIAL**

**Supplementary Figures**

**Figure S1 – SERBP1-associated proteins isolated via pulldown experiments in 293T cells.** 293T cells were transfected with plasmids expressing SBP-SERBP1 or pUltra-SERBP1(control). SERBP1-associated proteins were isolated via pull-down with streptavidin beads. Images show the aspect of protein gels used in the mass spectrometry analysis. Three control and three experimental samples were obtained using two different cell lysis procedures. Figure shows uncropped gel.

**Figure S2 – SERBP1 in translation and ribosome biogenesis. A)** SERBP1 knockdown in U251 and U343 cells decreased translation as determined by puromycin incorporation assay**.** Control (C) and siSERBP1 knockdown (KD). **B)** Localization of SERBP1 in nucleoli of HeLa cells as indicated by transfected mGreen-SERBP1 corroborates its participation in ribosome biogenesis. **C)** mGreen-SERBP1 co-localizes with endogenous FBL (red) in the nucleoli of HeLa cells.

**Figure S3 – SERBP1-associated helicases and their respective domains.** Listed are all helicases found among identified SERBP1 interactors and their respective helicase domains (1–3), information on binding to DNA, RNA (2) or G4s (4, 5) and expression correlation with SERBP1 in different datasets (6–8). Green label indicates the presence of respective characteristics; ND = no data; ns = not significant. Datasets used to prepare the figure and detailed analyses are in Tables S1, S4, S5 and S6.

**Figure S4 – PARP, PARylation and SERBP1 function. A)** SERBP1 shows increased nuclear localization and co-localization with PARP1 in U251 cells after treatment with H_2_O_2_. **B)** Effect of PARylation/PAR binding on SERBP1 protein interactions. 293T cells were transfected with SBP-SERBP1. The experimental group was treated with PJ34 10mM for 2 hours. SERBP1-associated proteins were recovered via pulldown with streptavidin beads. Western blot shows PARylated proteins detected in input and pulldown using a poly-ADP-ribose binding reagent. **C)** Effects of combined siSERBP1 knockdown and PARP inhibitor treatment on GBM cell viability**.** Reduction in proliferation displayed by U251 and U343 cells treated with only PJ34 (PARP inhibitor) or SERBP1 siRNA (partial SERBP1 KD) and combination. The combination treatment showed a synergistic effect as judged by the Combination Index (CI) (9). Data are shown as means ± standard deviation. Statistical significance was evaluated by Student’s t-test.

**Figure S5 – SERBP1 interactors, PARylation, PAR- and G4-binding.** **A)** Venn diagram showing that many SERBP1-associated proteins that get PARylated and/or bind PAR (10–12) also bind to G4s (4, 5). **B)** Prevalence of RRM (2) and RGG (13) domains in those SERBP1 interactors that bind both PAR and G4s. Datasets used to prepare the figure and detailed analyses are in Tables S1, S5 and S6.

**Figure S6. SERBP1 association with G3BP1 in pathological stress granules and glioblastoma cells. A)** Representative co-immunofluorescence of G3BP1 and SERBP1 in control and AD brain tissues. Merged channel is represented. DAPI was used to stain nuclei. Magnification 20x and white scale bar: 100 µm. Inset 1 (Ctr) and Inset 2 (AD) selected from merged channels are represented in zoomed images (white scale bar: 5 µm). **B)** Quantification of SERBP1 puncta localized in stress granules (G3BP1 granules) in Ctr versus AD (unpaired t-test p<0.001,****). Quantification of stress granule density (G3BP1 positive foci) Ctr versus AD (ordinary one-way ANOVA p=0.006,***) and total SERBP1 puncta comparing Ctr versus AD (ordinary one-way ANOVA p<0.001,****). Quantification has been performed using BZ-X Analyzer software (Keyence) analyzing 3 frontal cortex sections (selected randomly) from 3 control and 3 AD cases. **C)** SERBP1 co-localization with G3BP1 in U251 cells.

**Supplementary Tables**

**Table S1 – SERBP1-associated proteins identified by pulldown in 293T cells.** Mass spectrometry results of pulled-down proteins with respective counts in SBP-SERBP1 and control cells in two different experimental conditions. The summary sheet displays gene names and UniProt IDs (2) of identified SERBP1 interactors. Additional sheets show SERBP1-associated proteins present in the nucleus and nucleolus. Also included are complete results of ShinyGO (1) and Metascape (14) GO enrichment analyses of SERBP1 and all its identified associated proteins as well as comparisons of newly identified SERBP1 interactors with previous proximity-label (BIO-ID) studies (15, 16) and compilations from BioGRID (17).

**Table S2 – Comparison between snoRNAs bound by SERBP1 and PARP1.** snoRNAs and scaRNAs bound by SERBP1 according to RIP-Seq analysis (18) and comparisons to PARP1 bound snoRNAs identified CLIP-identified (19).

**Table S3 – BioGRID data for STM1, SERBP1 yeast homolog.** STM1’s protein interactors as curated by BioGRID (17) with their gene ontologies (1), human homologs as derived from WORMHOLE (20), and overlap with SERBP1-associated proteins.

**Table S4 – Protein domains and intrinsically disordered regions of SERBP1-associated proteins.** Analysis of protein domains present in SERBP1-associated proteins (2) with their respective counts in reference to the total human proteome (3). Additional sheets display the results of gene ontology analyses for top-represented domains (21). Also shown are derived and curated intrinsically disordered proteins associated with SERBP1 according to MobiDB (22).

**Table S5 – Gene expression correlation analysis.** Significant expression correlation values for SERBP1 and associated proteins in glioblastoma, neuroblastoma, pancreatic adenocarcinoma, bladder urothelial carcinoma, sarcoma, and normal brain datasets (6, 7). Additionally, a list of SERBP1-associated proteins displaying a similar expression profile during cortex development according to Cortecon (8). The last sheet shows the top SERBP1 interactors with the highest number of high expression correlation instances and their respective proteomics counts from our pulldown experiment.

**Table S6 – Characteristics of SERBP1-associated proteins.** SERBP1-associated proteins are organized according to the presence of RGG boxes (13), RRM motifs (2), G4 binding (4, 5), PAR binding (12), PARylation (10, 11) and overlap with SARS-CoV-2 (23) and PARP1 (24) interactomes. Additional sheets show gene ontology results (1) of SERBP1-associated proteins that 1) get PARylated and/or bind to PAR, 2) overlap with SARS-CoV-2 interactors, or 3) overlap with PARP1 interactors. Mutual SERBP1- and PARP1-associated factors identified in an rRNA transcription and processing screening (25) are highlighted on the last sheet.

**Table S7 – SERBP1 and drug sensitivity.** Results of cell viability screening in SERBP1-overexpressing vs. control U343 cells and information on the utilized drugs. Separate sheets list genes conferring high sensitivity to different PARP inhibitors (26) and SERBP1 interactors identified in this study.

**Table S8 – Impact of SERBP1 and hnRNPU knockdown on the splicing of U251 cells.** List of splicing events and respective genes affected in siSERBP1 knockdown vs. control and sihnRNPU knockdown vs. control U251 cells. Additional files include splicing events affected by SERBP1 knockdown with evidence of SERBP1 binding sites (CLIP sites) (18, 27–29) in the proximity (<100nt) of regulated splice sites and splicing events affected by both SERBP1 and hnRNPU knockdown in the same direction.

**Table S9 – Participation of SERBP1-associated proteins and snoRNAs in membraneless organelles.** Proteins associated with SERBP1 and their presence in stress granules, paraspeckles, Cajal bodies, nuclear specks, nucleoli, P-bodies (30) and Tau aggregates (31). UniProt ID mapping (2) was used to match gene names and UniProt IDs. SERBP1 interactors implicated in membraneless organelles were evaluated for prevalence of intrinsic disorder (22), G4 binding (4, 5), PARylation (10, 11), and PAR binding (12). snoRNAs found in a previous study to be enriched in Tau aggregates (32) that were also identified as SERBP1 targets according to RIP-Seq analysis (18).
