## Supplementary Figure 1 for "SERBP1 interacts with PARP1 and is present in PARylation-dependent protein complexes regulating splicing, cell division, and ribosome biogenesis"

### Slide 1
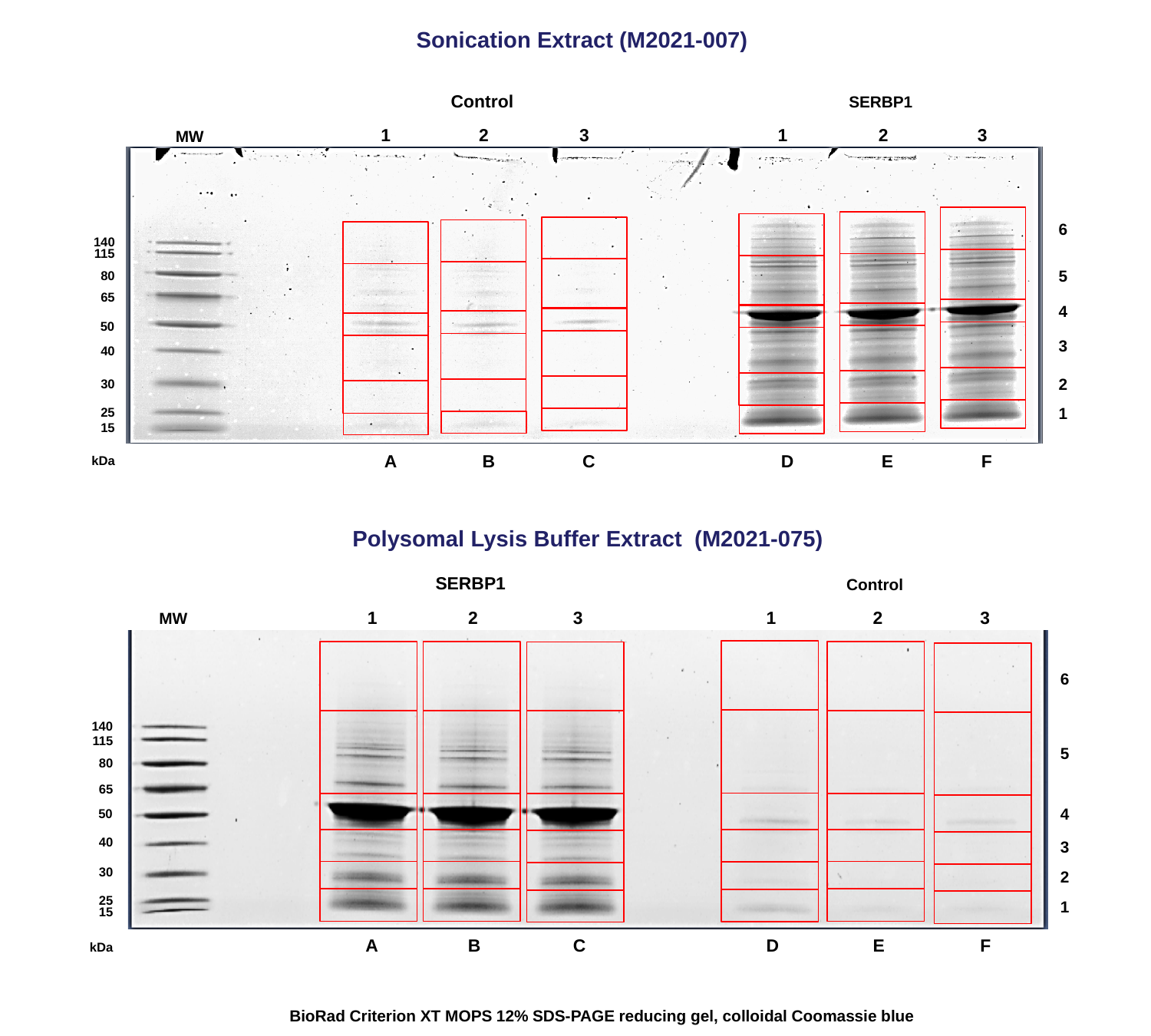

Sonication Extract (M2021-007)
Control
SERBP1
1
2
3
1
2
3
MW
6
5
4
3
2
1
140
115
80
65
50
40
30
25
15
A
B
C
D
E
F
kDa
Polysomal Lysis Buffer Extract (M2021-075)
SERBP1
Control
1
2
3
1
2
3
MW
6
140
115
5
80
65
4
50
40
3
30
2
25
1
15
A
B
C
D
E
F
kDa
BioRad Criterion XT MOPS 12% SDS-PAGE reducing gel, colloidal Coomassie blue
