## Supplementary Figure 2 for "SERBP1 interacts with PARP1 and is present in PARylation-dependent protein complexes regulating splicing, cell division, and ribosome biogenesis"

### Slide 1
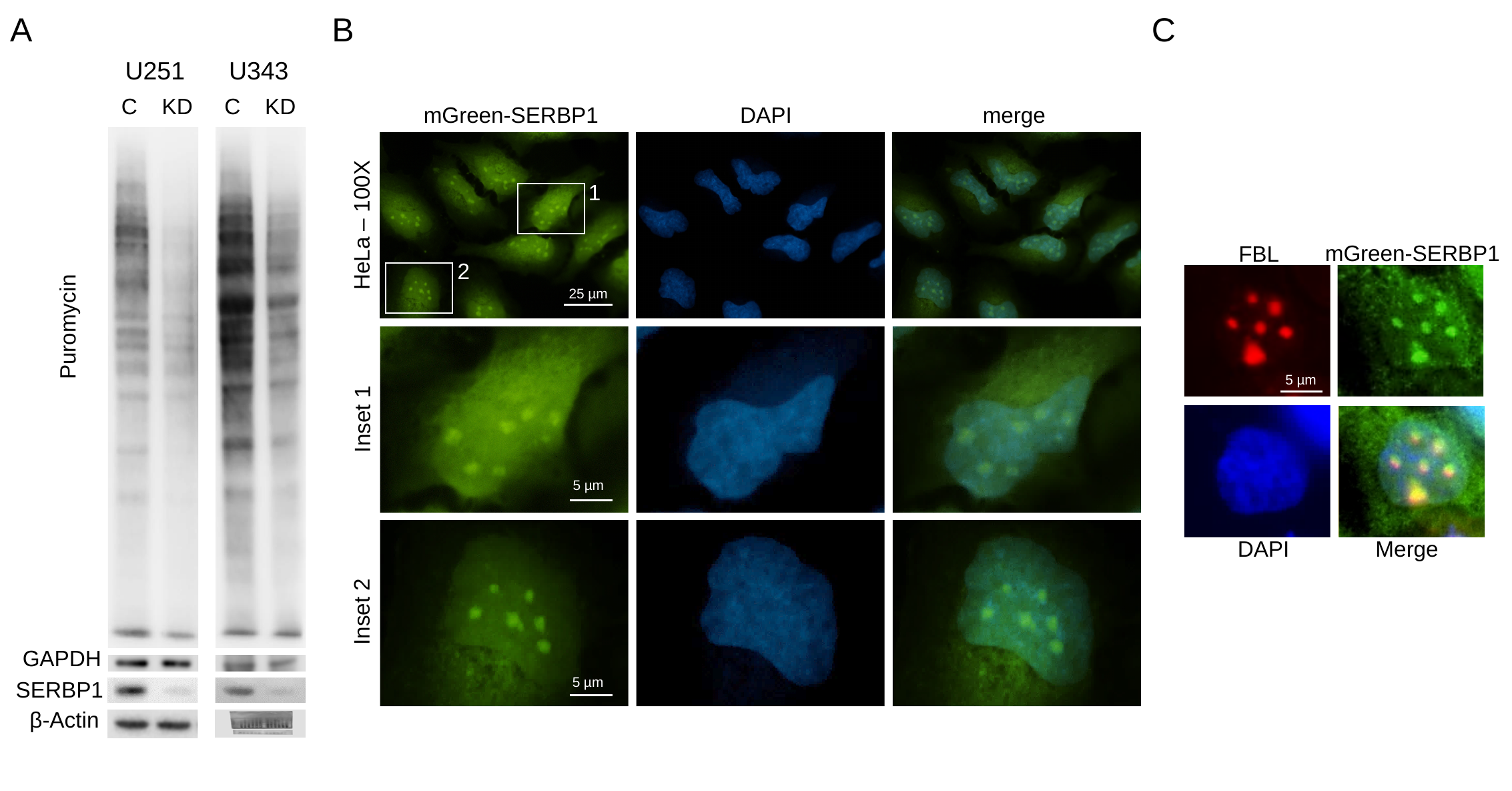

A
B
C
U251 U343
C KD
C KD
Puromycin
GAPDH
SERBP1
β-Actin
mGreen-SERBP1 DAPI merge
1
HeLa – 100X
mGreen-SERBP1
FBL
2
5 µm
25 µm
Inset 1
5 µm
DAPI Merge
Inset 2
5 µm
