## Supplementary figures and images for "SERBP1 interacts with PARP1 and is present in PARylation-dependent protein complexes regulating splicing, cell division, and ribosome biogenesis"

### Supplementary Figure 3

#### Slide 1
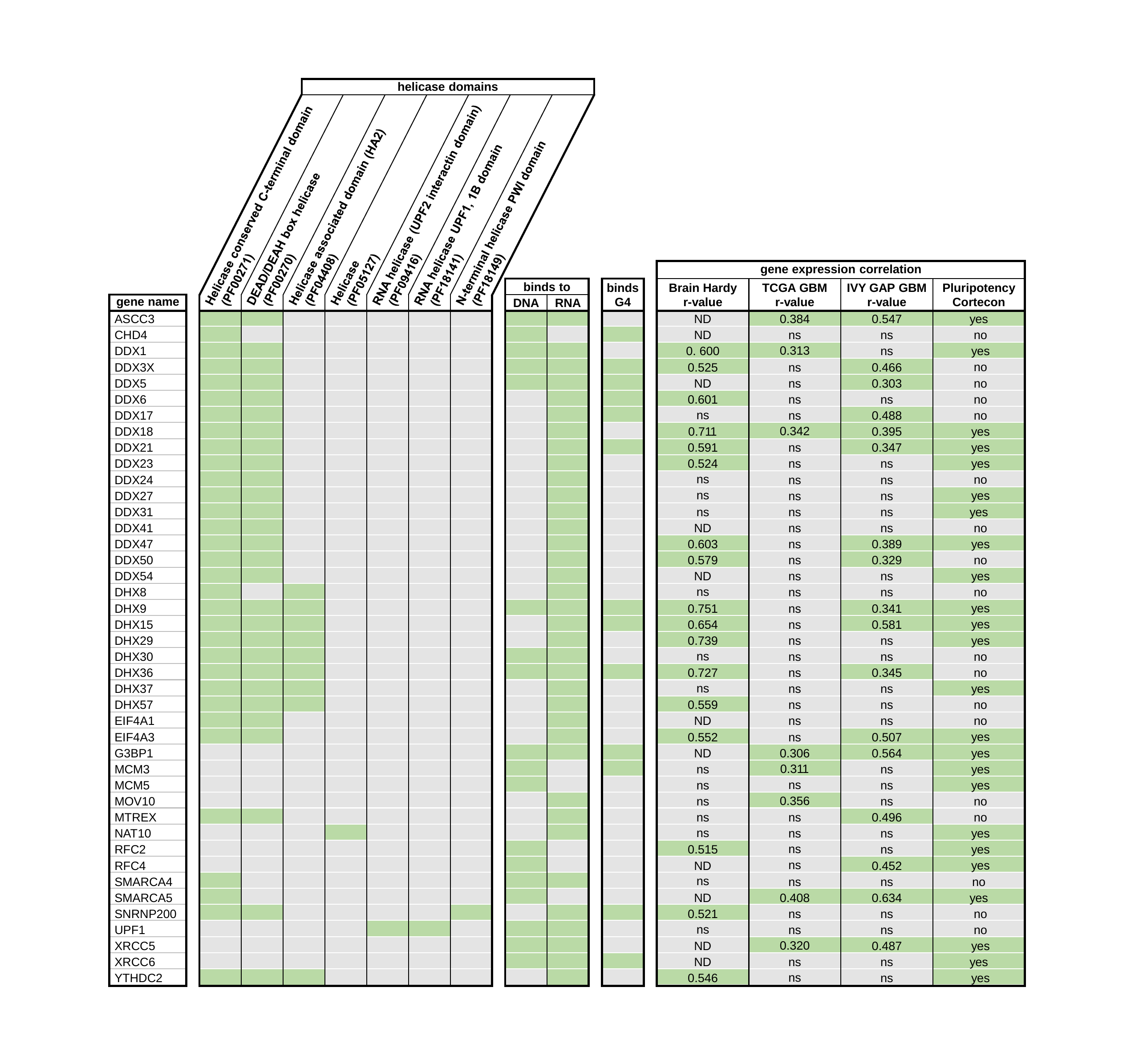

### Supplementary Figure 6

#### Slide 1
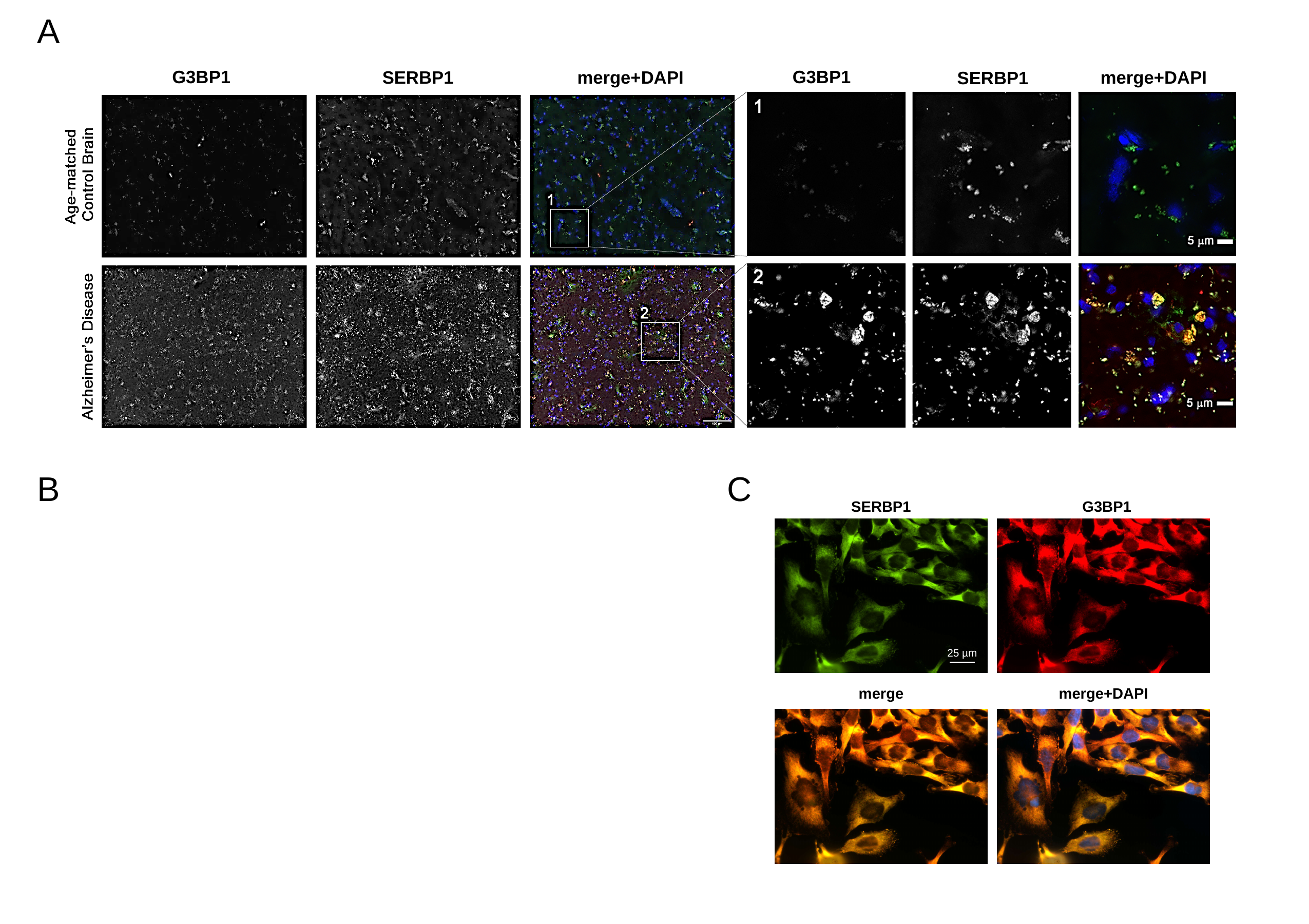

A
B
C
SERBP1
G3BP1
25 µm
merge
merge+DAPI
