## Supplementary Figure 4 for "SERBP1 interacts with PARP1 and is present in PARylation-dependent protein complexes regulating splicing, cell division, and ribosome biogenesis"

### Slide 1
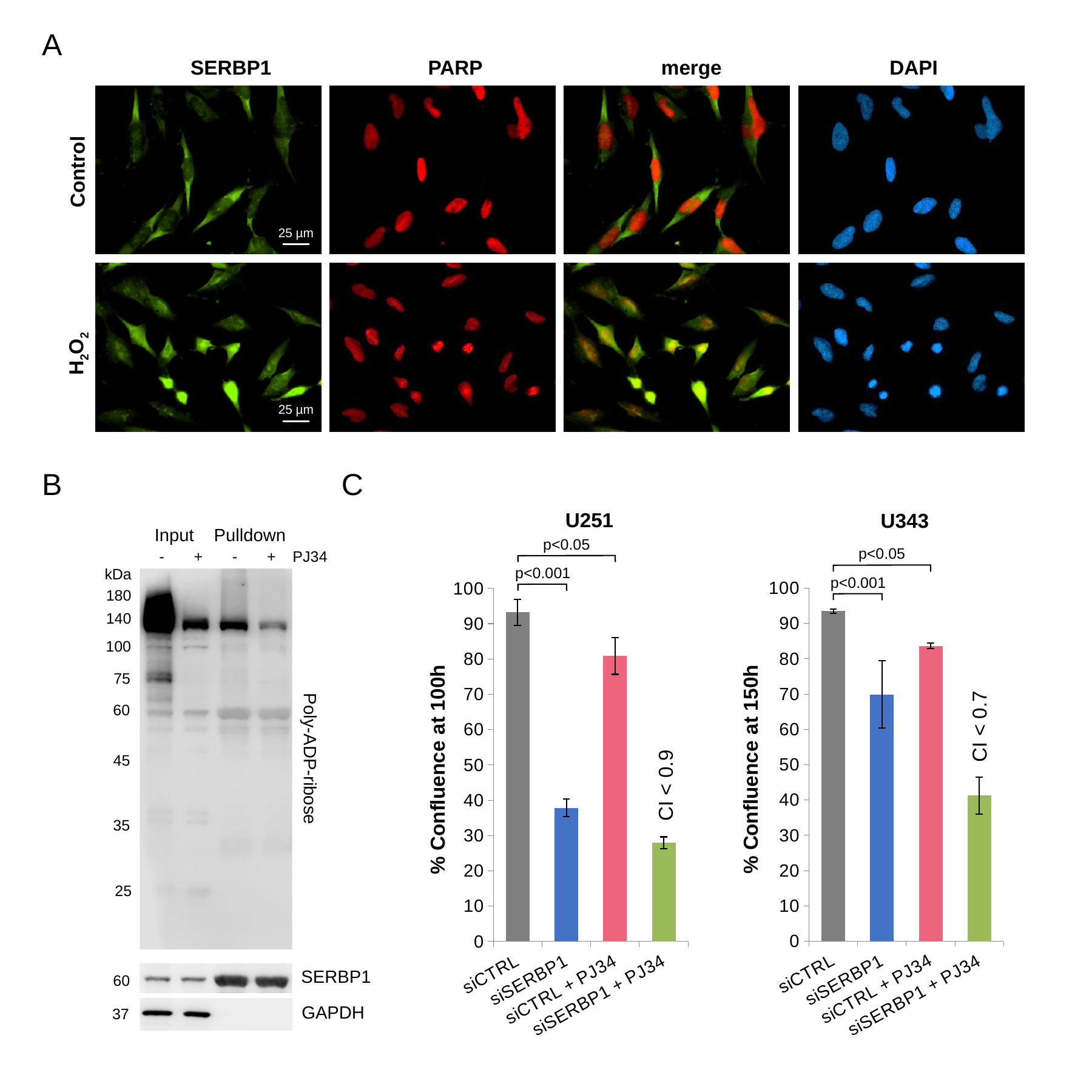

A
SERBP1 PARP merge DAPI
Control
25 µm
H_2_O_2_
25 µm
B
C
U251
U343
Input Pulldown
p<0.05
- + - + PJ34
p<0.05
kDa
p<0.001
#### Chart
| Category | |
|---|---|
| siCTRL | 93.47884 |
| siSERBP1 | 69.88965 |
| siCTRL + PJ34 | 83.66309 |
| siSERBP1 + PJ34 | 41.28157 |
#### Chart
| Category | |
|---|---|
| siCTRL | 93.17999999999999 |
| siSERBP1 | 37.845 |
| siCTRL + PJ34 | 80.88000000000001 |
| siSERBP1 + PJ34 | 27.945 |
p<0.001
180
140
100
75
60
CI < 0.7
Poly-ADP-ribose
45
CI < 0.9
35
25
SERBP1
60
GAPDH
37
