## Supplementary Figure 5 for "SERBP1 interacts with PARP1 and is present in PARylation-dependent protein complexes regulating splicing, cell division, and ribosome biogenesis"

### Slide 1
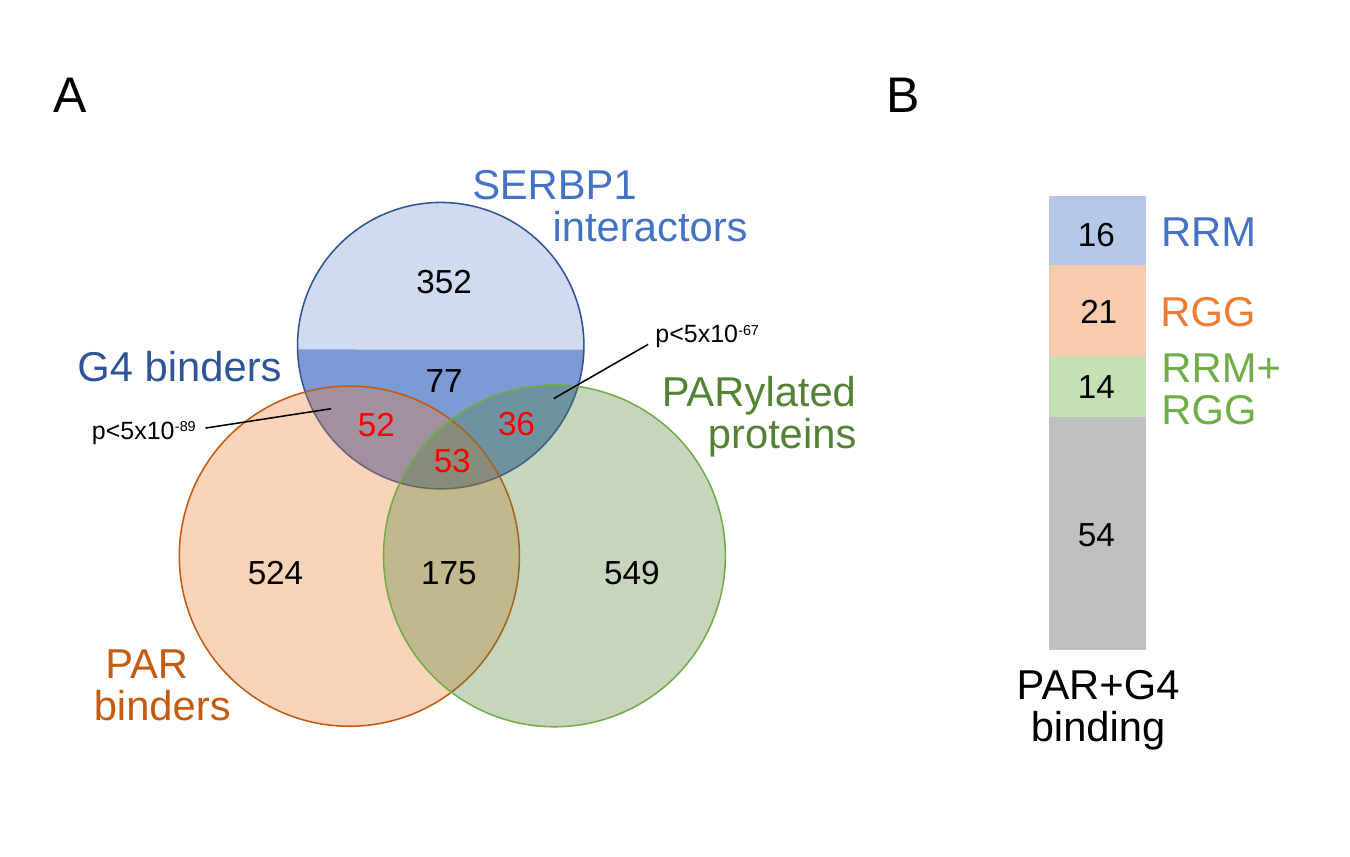

A
B
#### Chart
| Category | neither | both | RGG | RRM |
|---|---|---|---|---|
| PAR+G4 binders | 54.0 | 14.0 | 21.0 | 16.0 |16
RRM
21
RGG
RRM+RGG
14
54
PAR+G4 binding
SERBP1
 interactors
352
p<5x10^-67^
G4 binders
77
PARylated
 proteins
36
52
p<5x10^-89^
53
524
175
549
 PAR
binders
