## Supplementary material for "SERBP1 interacts with PARP1 and is present in PARylation-dependent protein complexes regulating splicing, cell division, and ribosome biogenesis": List of key reagents

| **Key Resources Table** | | | | |
| --- | --- | --- | --- | --- |
| **Reagent type (species) or resource** | **Designation** | **Source or reference** | **Identifiers** | **Additional information** |
| gene (*Homo sapiens*) | SERBP1 | GenBank | HGNC:HGNC:17860 |  |
| cell line (*Homo sapiens*) | U251 (glioblastoma) | Uppsala University |  |  |
| cell line (*Homo sapiens*) | U343 (glioblastoma) | Uppsala University |  |  |
| cell line (*Homo sapiens*) | 293T (normal) | ATCC |  |  |
| transfected construct (human) | SERBP1 SMARTpool siRNA | Dharmacon | Cat#: L-020528-01-0005 |  |
| biological sample (human) | Control and Alzheimer’s Disease (AD) human brains | Institute for Brain Aging and Dementia at UC Irvine |  | Braak Stage V-VI for AD brains |
| antibody | anti-SERBP1 (mouse monoclonal) | Santa Cruz Biotechnology | Cat#:sc100800 | WB (1:100 for human, 1:1000 for cell line samples), IF (1:50 for human, 1:200 for cell line samples), PLA (1:100) |
| antibody | anti-GAPDH (rabbit polyclonal) | Abcam | Cat#: ab9485 | WB (1:1000) |
| antibody | anti-G3BP1 (rabbit polyclonal) | Cell Signaling | Cat#:17798S | IF (1:200) |
| antibody | anti-PARP1 (rabbit monoclonal) | Cell Signaling | Cat#:9532S | WB (1:1000), PLA (1:200) |
| antibody | anti-SYNCRIP (rabbit polyclonal) | Invitrogen | Cat#:PA5-59501 | WB (1:1000) |
| antibody | anti-hnRNPU (rabbit monoclonal) | Cell Signaling | Cat#:34095 | WB (1:1000) |
| antibody | anti-Puromycin (mouse monoclonal) | Kerafast | Cat# EQ0001 | WB (1:1000) |
| antibody | anti-His (mouse monoclonal) | Santa Cruz Biotechnology | Cat#:sc-53073 | WB (1:1000) |
| antibody | anti-Flag (mouse monoclonal) | Invitrogen | Cat#:MA1-91878 | WB (1:1000) |
| antibody | anti-GAPDH (mouse monoclonal) | Santa Cruz Biotechnology | Cat#:SC32233 | WB (1:2000) |
| antibody | anti-β-Actin (rabbit polyclonal) | Abcam | Cat#: ab8227 | WB (1:1000) |
| antibody | anti-β-Tubulin (mouse monoclonal) | Sigma-Aldrich | Cat#:T8328 | WB (1:2000) |
| antibody | anti-α-Tubulin (mouse monoclonal) | Invitrogen | Cat#:236-10501 | IF (1:200) |
| antibody | anti-FBL (rabbit monoclonal) | Cell Signaling | Cat#:C13C3 | WB (1:1000) |
| antibody | anti-NCL (rabbit monoclonal) | Cell Signaling | Cat#: D4C70 | WB (1:1000) |
| antibody | anti-G3BP1 (mouse monoclonal) | Abcam | Cat#: ab56574 | IF (1:200) |
| antibody | Anti-phospho-Tau (Thr231) (mouse monoclonal) | Thermo Fisher | Cat#:MN1040 | IF(1:250), PLA(1:500) |
| antibody | HRP-conjugated anti-rabbit (goat polyconal) | Santa Cruz Biotechnology | Cat#:sc-2030 | WB (1:5000) |
| antibody | HRP-conjugated anti-mouse (goat, polyclonal) | Santa Cruz Biotechnology | Cat#:sc-2005 | WB (1:5000) |
| antibody | Alexa Fluor 488-conjugated anti-rabbit (goat polyclonal) | Invitrogen | Cat# A11008 | IF (1:500) |
| antibody | Alexa Fluor 568-conjugated anti-mouse (goat polyclonal) | Invitrogen | Cat#: A11004 | IF (1:500) |
| antibody | Anti-poly-ADP-ribose binding reagent (with rabbit Fc-tag) | Millipore | Cat#: MABE1031; RRID: AB_2665467 | WB (1:1000) |
| recombinant DNA reagent | pEF1 (plasmid) | Thermo Fisher | Cat#: V92120 |  |
| recombinant DNA reagent | pSBP-SERBP1 (plasmid) | This paper |  | SERBP1 ORF and SBP-tag cloned in frame in pEF1 backbone |
| recombinant DNA reagent | pUltra-SERBP1 lentiviral vector | DOI:10.1186/s13059-020-02115-y |  | control for pulldown experiments (no SBP-tag) |
| recombinant DNA reagent | pcDNA3.1-mGreenLantern (plasmid) | Addgene | Plasmid:#161912 |  |
| recombinant DNA reagent | mGreen-SERBP1 (plasmid) | This paper |  | SERBP1 ORF cloned in frame in pcDNA3.1-mGreenLantern backbone |
| recombinant DNA reagent | Flag-PARP1 | Addgene | Plasmid:#111575 |  |
| peptide, recombinant protein | 6xHis-tagged SERBP1 | DOI: 10.3389/fmolb.2021.744707 |  |  |
| commercial assay or kit | Lipofectamine RNAiMAX | Invitrogen | Cat#:13778150 |  |
| commercial assay or kit | Streptavidin beads | GE Healthcare Life Sciences | Cat#:17-5113-01 |  |
| commercial assay or kit | Cell Titer Glo 2.0 | Promega | Cat#: G9243 |  |
| commercial assay or kit | Duolink® PLA in Situ Red starter kit mouse/rabbit | Sigma-Aldrich | Cat#: DUO92101 |  |
| chemical compound, drug | Cambridge Cancer Compound Library | Selleck Chem | Cat#: L2300 | 100 nM in 0.1% DMSO treatment concentration |
| chemical compound, drug | Puromycin | Sigma-Aldrich | Cat#: P7255 |  |
| chemical compound, drug | Paclitaxel | Cayman Chem | Cat#:10461 |  |
| chemical compound, drug | PARP inhibitor PJ34 | Enzo | Cat#: ALX-270-289 |  |
| software, algorithm | Mascot v2.7.0 | Matrix Science |  |  |
| software, algorithm | Scaffold v4.9.0 | Proteome Software |  |  |
| software, algorithm | ImageJ FIJI | NIH |  |  |
| software, algorithm | BioInfoRx | URL: https://bioinforx.com/apps/venn.php |  |  |
| software, algorithm | Nematode genome comparison browser | URL: http://nemates.org/MA/progs/overlap_stats.html |  |  |
| software, algorithm | ShinyGO v0.67 & v0.77 | DOI:10.1093/bioinformatics/btz931 |  |  |
| software, algorithm | Metascape v3.5 | DOI:10.1038/s41467-019-09234-6 |  |  |
| software, algorithm | Panther v17.0 | DOI:10.1002/pro.4218 |  |  |
| software, algorithm | Revigo | DOI:10.1371/journal.pone.0021800 |  |  |
| software, algorithm | STRING v11.5 | DOI:10.1093/nar/gkac1000 |  |  |
| software, algorithm | Cytoscape | DOI:10.1021/acs.jproteome.2c00651 |  |  |
| software, algorithm | R2 | URL:http://r2.amc.nl |  |  |
| software, algorithm | dcGO Enrichment mining service | DOI:10.1093/nar/gks1080 |  |  |
| software, algorithm | STAR v2.7.7.a | DOI:10.1093/bioinformatics/bts635 |  |  |
| software, algorithm | rMATS v4.1.2 | DOI:10.1073/pnas.1419161111 |  |  |
| software, algorithm | rmats2sashimiplot tool | https://github.com/Xinglab/rmats2sashimiplot |  |  |
| software, algorithm | BEDTools intersect software | DOI:10.1093/bioinformatics/btq033 |  |  |
| software, algorithm | Kallisto v0.46.1 | DOI:10.1038/nbt.3519 |  |  |
| software, algorithm | R package tximport | DOI:10.12688/f1000research.7563.1 |  |  |
| software, algorithm | DESeq2 | DOI:10.1186/s13059-014-0550-8 |  |  |
| other | Fluor Save | Invitrogen | Cat#:345789 |  |
| other | Prolong Gold Antifade with DAPI | Thermo Fisher | Cat#: P36931 |  |
